# Microsnoop: A Generalized Tool for Unbiased Representation of Diverse Microscopy Images

**DOI:** 10.1101/2023.02.25.530004

**Authors:** Dejin Xun, Rui Wang, Xingcai Zhang, Yi Wang

**Author notes:** Corresponding author. (Y.W.); (X.C.Z.); (R.W.).

## Abstract

Microscopy image profiling is becoming increasingly important in biological research. Microsnoop is a new deep learning-based representation tool that has been trained on large-scale microscopy images using masked self-supervised learning, eliminating the need for manual annotation. Microsnoop can unbiasedly profile a wide range of complex and heterogeneous images, including single-cell, fully imaged, and batch-experiment data. Its performance was evaluated on seven high-quality datasets, containing over 358,000 images and 1,270,000 single cells with varying resolutions and channels from cellular organelles to tissues. The results show that Microsnoop outperforms previous generalist and even custom algorithms, demonstrating its robustness and state-of-the-art performance in all biological applications. Furthermore, Microsnoop can contribute to multi-modal studies and is highly inclusive of GPU and CPU capabilities. It can be easily and freely deployed on local or cloud computing platforms.

## Introduction

Automatic quantitative profiling of microscopy images has become increasingly ubiquitous in a broad range of biological research, spanning from small-scale investigations to high throughput experiments^1^. The analysis of visual phenotypes, which involves profiling intricate information from images, has demonstrated its usefulness in diverse areas of biology^2^. These include protein localization^3^, cell cycle stage classification^4^, mechanisms of action predictions^5^, and high-content drug discovery^6^. Additionally, the emergence of spatial omics has given rise to new requirements for the quantification of microscopy images. For example, spatial proteomics methods can image more than 50 disease-related proteins in a single tissue slice^7^, while spatial transcriptomics allows for the simultaneous acquisition of both image data and transcriptional profiles^8^. These developments underscore the need for a high-performance, generalist representation tool that can effectively handle heterogeneous microscopy images.

The traditional approach to profiling microscopy images involves extracting predefined morphological features, such as intensity, shape, texture, granularity, and colocalization^9–10^. However, this method has several limitations, including low computational efficiency, potential information loss, and sensitivity to image quality^11^. To overcome these deficiencies, recent advancements in computer vision and deep learning have given rise to learning-based feature extraction methods that use representation learning. This technique involves pre-training a model on pretext tasks and then using part of the network as a feature extractor for downstream analysis. These methods can be divided into two categories: task-oriented custom methods and generalist methods. Task-oriented methods^4, 12–15^ are pre-trained on data from the same source and designed specifically for biological research, such as cell cycle stage prediction. In contrast, generalist methods require training data that are not specific to any particular biological problem. One of the most widely used generalist methods involves using models trained for ImageNet^16^ (a natural image classification task), which has also been utilized in recent multi-modal research^17^.

However, the extent to which the feature extraction patterns learned from natural images can capture the subtle phenotypes of microscopy images has not been fully validated by comparative research. To better match the feature domain to downstream microscopy image profiling tasks, the CytolmageNet^18^ study was conducted, where image representation was learned based on a microscopy image classification task (890K images,894 classes). Although this study demonstrated comparable performance to ImageNet, it still relied on the supervised learning approach that can be labor-intensive, prone to biases from semantic annotations, and potentially increase the difficulty of achieving higher representation performance.

The field of microscopy image analysis can greatly benefit from the development of an unbiased, high-performance, generalist image representation tool. Beyond facilitating accurate downstream analysis, such a tool would enable unsupervised analysis for identifying new phenotypes. It can facilitate the separation of feature extraction and downstream analysis process, allowing for downstream analysis conducted on computers with limited computing power. The representations of images that are much smaller than the original images can be easily stored and transferred, and private data can be shared securely through these representations without disclosing the original images. In addition, secondary analysis becomes possible, such as the creation of large image databases or joint analysis with other data representations. Nevertheless, the complexity and diversity of microscopy images pose significant challenges in the development of such a tool.

Self-supervised representation learning offers a promising solution by allowing the model to learn directly from pixels without relying on pre-defined semantic annotations. This approach involves transforming the original images and training the model to learn the mapping between the transformed and original image. Various transformation methods have been employed, such as direct copying^19^, partial channel drop^20^, or image masking^21^, with masked visual representation learning being particularly popular in natural image studies^22–24^. Recent advancements in cell segmentation algorithms have also indicated the remarkable generalization ability of networks trained on generalized data^25–27^. However, developing a universal tool for microscopy image profiling presents several challenges, including handling images with varying resolutions and channel numbers (such as 1, 2, 3, 5 and 56)^3–4, 7, 26, 28^, joint representation learning for multiple image styles, processing various image types, and addressing technical variations in high-content experiments that may introduce batch effects in the feature space^29–30^.

This study presents Microsnoop, a universal tool for the impartial representation of microscopy images using masked self-supervised learning. The proposed pipeline is capable of handling heterogeneous images and includes a task distribution module to cater to users with varying computing power. To meet diverse image profiling requirements, the images are categorized into three types with corresponding pipelines. The performance of Microsnoop was assessed using seven evaluation datasets from various biological studies and compared against both generalist and custom algorithms. The findings demonstrate Microsnoop’s robust feature extraction capabilities and potential for analyzing multi-modal biological data. The tool is freely available at https://github.com/cellimnet/microsnoop-publish.

## Results

### The design of a generalist representation tool

In this study, we developed a generalist tool called Microsnoop for the unbiased representation of microscopy images through masked self-supervised learning. As large and diverse datasets are beneficial for the training of generalist models, we collected and curated 10,458 high-quality microscopy images from various sources published by the cell segmentation community^25–27, 31–33^. These images were taken using different technologies and have different resolutions and channel numbers, with channels ranging from cellular organelles to tissues. The four main types of images include fluorescence, phase-contrast, tissue and histopathology images (Fig. 1a(i) and Supplementary Table 1). To accommodate the variable number of image channels, the input to the neural network was set as one-channel images (related to one-channel feature concatenation strategy below). All images channels in the training set were split out and further selected to form a one-channel data pool (Methods). Before training, images in each batch were preprocessed in three steps: (1) Sample: randomly select one batch of images from the four types in turn to reduce the effects of unequal amounts of data; (2) Augment: randomly crop a 224*224 region (pad if smaller) from each image, then normalize, random rotate and scale the image, with the result serving as the network target; (3) Transform: randomly mask a portion of the target image patches, with the result serving as the network input. In terms of network architecture design, this study employed a CNN-based^34^ (convolution neural network) architecture, despite the growing interest in Transformer-based architectures^35^ for natural image analysis. This choice was motivated by the superior performance observed for the CNN architecture in our preliminary evaluations (Extended Data Fig. 1 and Methods). This performance disparity may be attributed to the difference in the amount of training data provided. Typically, the pre-training of a ViT architecture^36^ requires a large corpus of data, with over 1 million or even 1 billion images used in the case of natural image studies^21^. However, our microscopy image dataset involved a relatively smaller set of training data, which may not have been sufficient to provide adequate training for the Transformer-based architecture.

**Fig. 1.**
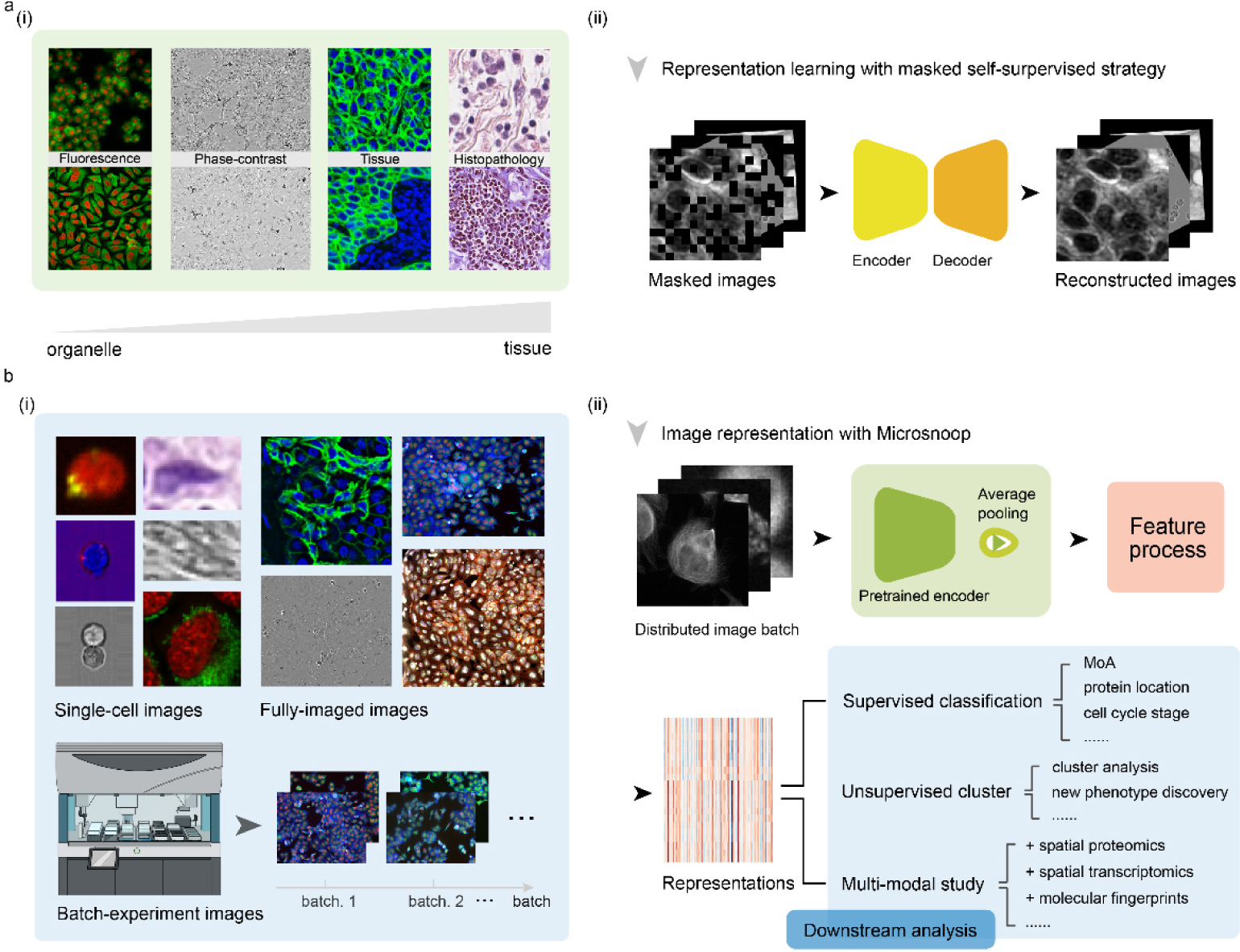
Design of Microsnoop for microscopy image representation. **a,** Schematic of the learning process. (i) Example of the four main category images are shown. The channels range from cellular organelles to tissues. (ii) A masked self-supervised learning strategy was employed and only images are required for training without additional manual annotation. One-channel masked images were set as the input and the Encoder-Decoder were required to reconstruct the original images. **b,** At test time, (i) Example images from various downstream tasks are shown, with different resolutions, number of channels and image types. These microscopy images are categorized into 3 types to ensure the broad coverage of image profiling needs. (ii) Application of Microsnoop. Firstly, images are managed by an in-built task distribution module (Fig. 3a), which generates one batch one-channel images for feature extraction. Each batch of images is fed into the pre-trained encoder, and the output smallest convolutional maps are processed by average pooling. Then, all extracted embeddings are processed according to different profiling tasks (introduced in the following section). The potential downstream analyses of our generalist representation tool are shown in the panel.

We employed a masked self-supervised learning strategy to train the network, where a randomly selected percentage of image patches are masked and used as inputs. The network was then tasked with reconstructing the original, unmasked images. During training, masked images are encoded into high-level features through four consecutive downsampling steps, and the process of image reconstruction is accomplished through mirror-symmetric upsampling (Fig. 1a(ii)). The learning process is guided by minimizing the self-supervision loss function (Methods), which promotes the model to learn useful features that enable it to recover the masked parts of the images based on the information present in the remaining parts. This is a challenging task, which necessitates a comprehensive understanding that transcends simple low-level image statistics.

At test time, a generalist tool needs to face a range of image processing needs. To cater for this condition, we chose to categorize images based on the image profiling process itself, rather than solely on their biological applications that may be limited in scope. Our categorization comprises three types: single-cell images, fully-imaged images, and batch-experiment images. (Fig. 1b(i)). The images to be processed are first managed by an in-built task distribution module (below), and then fed into the pre-trained encoder on a batch-by-batch basis for feature extraction. The output smallest convolutional maps are processed through global average pooling to produce initial 256-dimensional feature embeddings. Subsequently, feature aggregation is performed in accordance with different profiling tasks (details provided below). The final image representations can be used for various downstream analyses (Fig. 1b(ii)).

### Diversified evaluation datasets

In prior studies, attention was primarily focused on a limited number of specific datasets^5, 37–39^. In our work, to give a more comprehensive evaluation of our generalist tool, we collected and curated 7 evaluation datasets, encompassing commonly used datasets along with some novel additions, comprising over 358,000 images and 1,270,000 single cells (Methods and Extended Data Fig. 2). These images showcase a diverse array of characteristics, including various resolutions, image types, number of channels, and biological applications, such as protein localization estimation, cell cycle stage identification, and MoA prediction (Supplementary Table 2). In our study, four of the seven evaluation datasets focused on single-cell images. The performance of the model on fluorescent images, including bright-field channels, was assessed by COOS7 Test 1-4^39^, CYCLoPs^3^ and BBBC048^4^. For the assessment of the model’s ability to handle more challenging histopathology images, we employed the CoNSeP^40^ dataset. The LIVECell Test^26^ and TissueNet Test^27^ datasets were designed to evaluate a model’s performance on fully-imaged image classification tasks, involving phase-contrast and tissue image representation, respectively. Lastly, the BBBC021^41^ dataset was employed to evaluate the representation ability of the model for batch-experiment images.

### Microsnoop accurately reconstructs the masked input images

In the investigation of optimal mask ratio for learning features from microscopy images, we found that a 25% mask was optimal for this task. This was determined by testing 8 different mask ratios (5%, 15%, 25%, 35%, 45%, 55%, 65% and 75%) and comparing the results (Extended Data Fig. 3). To get a qualitative sense of the reconstruction task, we showed an example of each image type from the validation set (Fig. 2a). By inputting the 25% masked image into the pre-trained network, we were able to produce a reconstructed image that closely resembles the original, with only some detailed textures lost. This level of detail recovery is not easily achievable by humans. The reconstruction results on single-cell images from the evaluation datasets were even more impressive, with the reconstructed image being nearly indistinguishable from the original image (Fig. 2b and Extended Data Fig. 4). The improved performance on single-cell images in comparison to fully-imaged ones can be attributed to cellular heterogeneity, which results in diverse cell phenotypes. The abundance of reference information from single-cell images allows for the more successful recovery of a limited number of instances. These results demonstrate that the pre-trained Microsnoop network, has learned good representations of the microscopy images.

**Fig. 2.**
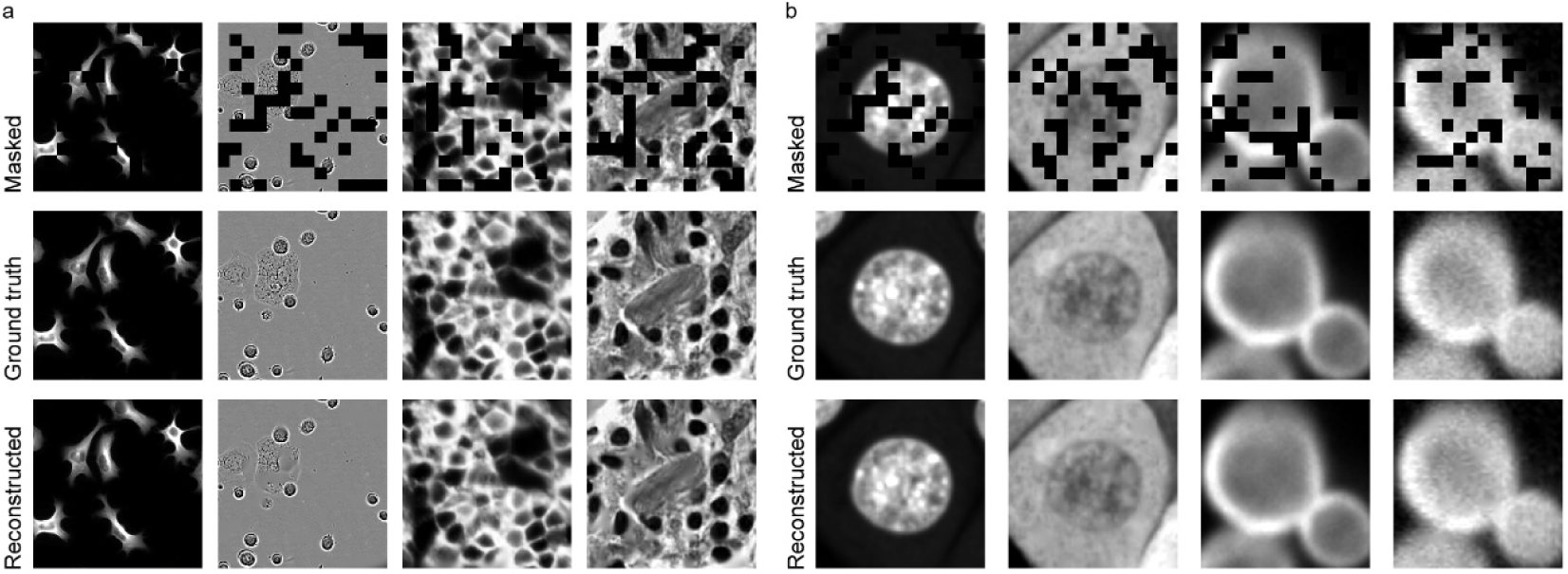
Reconstruction results with Microsnoop. **a,** Example results for images from the validation set, with a masking ratio of 25% applied on inputs. One representative image is selected for each image type. **b,** Example results for single-cell images from evaluated data, with a masking ratio of 25% applied on inputs. The left two columns are from COOS7 and the right two are from CYCLoPs. Two representative images (different imaging channels of the same cell) are selected for each dataset. Example results on other evaluated datasets are shown in Extended Data Figs. 4.

### Microsnoop profile of single-cell images with one-channel feature concatenation

Single-cell images can be produced directly by an imaging instrument such as imaging flow cytometry (IFC)^42^, or obtained through cell segmentation processing on fully-imaged images. To accommodate the variable number of channels, we devised a one-channel feature concatenation strategy (Fig. 3a). Each channel of the multi-channel image is processed independently by Microsnoop, and the resulting embeddings are concatenated in an orderly manner. To prevent confusion during processing, a unique index is assigned to each image when multiple images are being processed. To address potential memory overflow issues when processing large batches of data, we established a task distribution module. This module efficiently manages image pathways and distributes images for processing, read into the CPU and transferred to the GPU as needed. The user is empowered to optimize performance by adjusting parameters according to the available memory capacity of both the CPU and GPU. Furthermore, our system features a scalable, distributed design, which is capable of supporting multiple GPUs, providing a solution for increasing data demands.

**Fig. 3.**
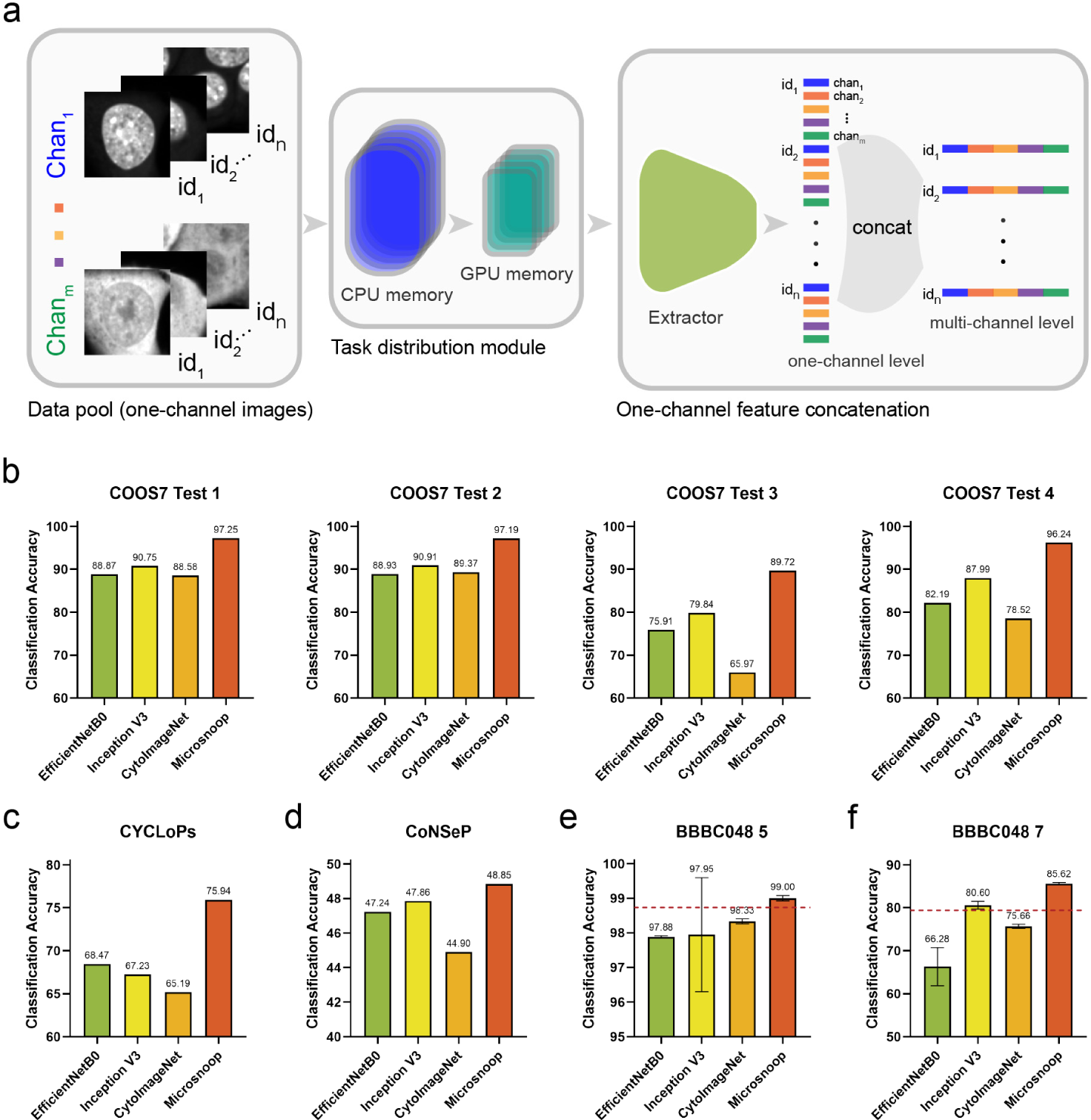
Profiling with Microsnoop on single-cell images. **a,** Pipeline. Every channel of the single-cell image is processed independently, and the one-channel level embeddings are concatenated to get multi-channel level image representations. A task distribution module is provided to prevent memory overflow. The Extractor denotes the pretrained encoder combined with the average pooling layer shown in Fig. 1a(ii). **b-f,** Benchmarks. **b,** Benchmark on COOS7, containing four separate test sets. **c,** Benchmark on CYCLoPs. **d,** Benchmark on CoNSeP. **e,f,** Benchmarks on BBBC048, with two different classification tasks. Performances reported by the original paper are shown with dotted red lines. Error bars represent the mean ± SD of fivefold cross-validation results.

In our benchmark, we included three previous developed generalist methods in the comparisons: EfficientNetB0^43^, Inception V3^44^, CytoImageNet^18^, and custom methods that are accessible (Methods). For the COOS7 Test 1-4, CYCLoPs and CoNSeP, we evaluated performance with the K-Nearest Neighbor (KNN) classification accuracy (match between prediction and ground truth using the KNN classifier, which has been utilized in prior study^18^). For the dataset BBBC048, we used fivefold cross-validation for dataset split and evaluated the performance with the multilayer perceptron (MLP) classification accuracy (match between prediction and ground truth using the MLP classifier, as employed in the original paper^4^). Our evaluations revealed the exceptional performance of Microsnoop, which consistently outperformed all other methods. In the majority of cases, Microsnoop achieved significant improvements of more than 6%, and up to 10% (Fig. 3b-f). Notably, for the 7-classification task of BBBC048, Microsnoop achieved an accuracy of 85.62% without using any data from the dataset, surpassing the custom supervised learning algorithm in the original paper by 5.02%.

### Microsnoop profile of fully-imaged images with cell region cropping

Fully-imaged images are a common format directly obtained from most microscopes. Cell segmentation is usually the first step of phenotype profiling due to the inherent heterogeneity of cells. Although various generalist segmentation algorithms^25–27^ have been developed along with some fine-tuning strategies^45–46^, they may still introduce unwanted segmentation errors. For instance, in a large drug screening experiment, cell body images can present a range of phenotypes, and a segmentation algorithm may perform well on some but poorly on others (Extended Data Fig. 5a), potentially leading to unpredictable impacts on downstream analysis. To mitigate these issues, we introduced a cell region cropping strategy, where the segmentation algorithm is applied only on the easiest channel, such as the nucleus channel, which presents more robust segmentation results (Extended Data Fig. 5b). Cell regions are computed and cropped based on the segmentation masks and rescale constant (Fig. 4a(i) and Methods). Then, Microsnoop extracts features from the cropped single-cell images as described above (Fig. 4a(ii)). Finally, the resulting single-cell level embeddings are aggregated by computing their mean to obtain the fully-imaged level representations (Fig. 4a(iii)).

**Fig. 4.**
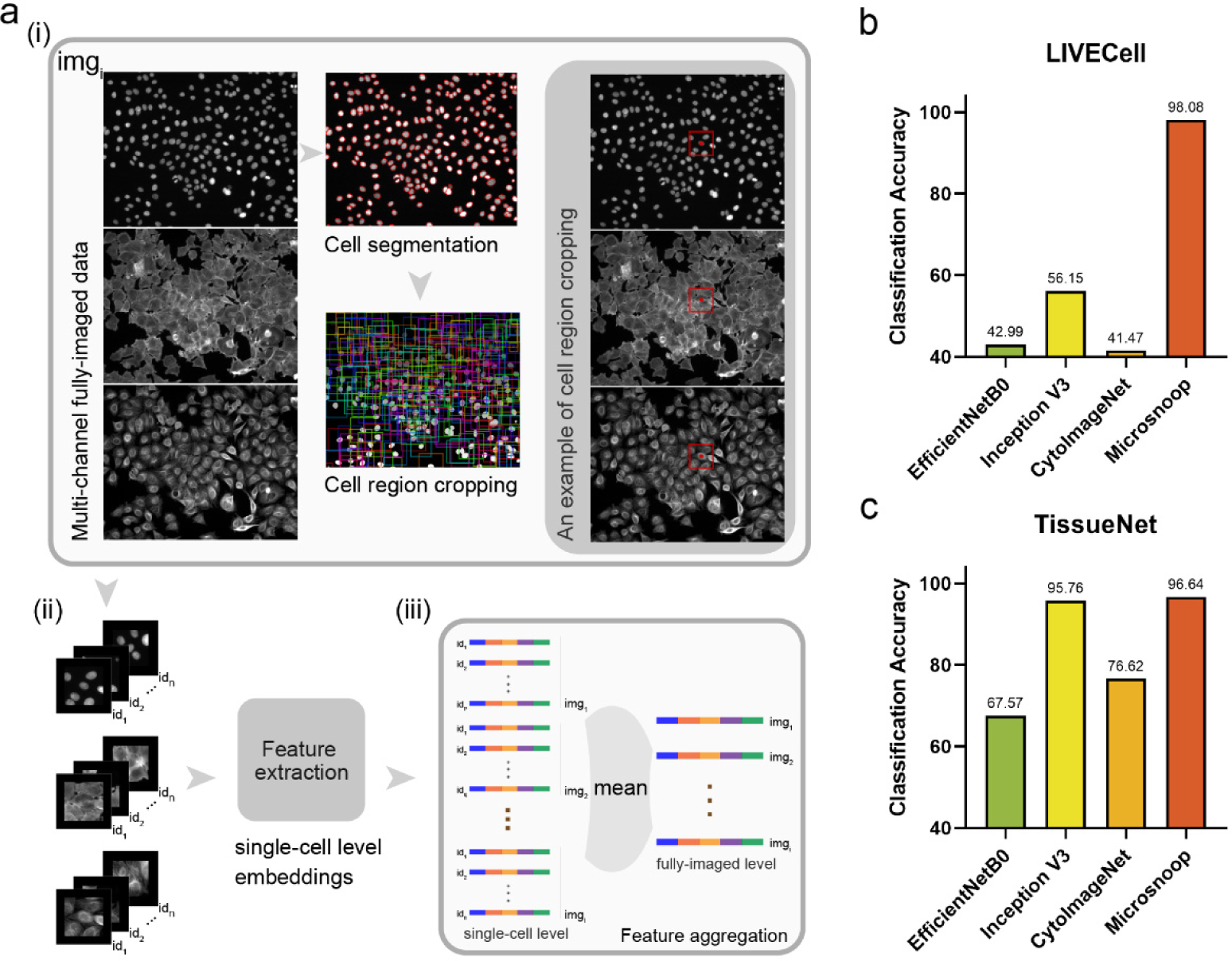
Profiling with Microsnoop on fully-imaged images. **a,** Pipeline. (i) Cell segmentation algorithm is conducted on the easiest channel (such as the nucleus channel) of the multi-channel fully-imaged image, then the cell region for each single cell is computed and cropped. (ii) Multi-channel single-cell images are processed as Fig. 3a, and (iii) the output single-cell level embeddings are aggregated to obtain the fully-imaged level image representations. **b,** Benchmark on LIVECell. **c,** Benchmark on TissueNet.

We evaluated the representation ability of Microsnoop on two fully-imaged image phenotype classification tasks, and tested previously mentioned generalist algorithms for comparison. Both tasks were evaluated using the KNN classification accuracy. The results showed that Microsnoop again outperformed other methods, and even a 41.93% improvement was obtained on the LIVECell Test dataset (Fig. 4b-c). Furthermore, Microsnoop showed strong inclusiveness to various image styles, with an accuracy of 98.08% on the LIVECell Test dataset and 96.64% on TissueNet Test.

### Microsnoop profile of batch-experiment images with sphering batch correction

In high-content screening experiments, batch effects due to technical variability can affect downstream analysis^29–30, 37–38^ (Fig. 5a). To address this issue, we employed a sphering batch correction method^47^. This assumes that the large variations observed in negative controls are associated with confounders, and any variation that is not observed in controls is associated with phenotypes. Sphering transformation aims to separate phenotypic variation from confounders. In our image representation pipeline for batch-experiment images, Microsnoop first extracts features from the fully-imaged images (as described above), and the resulting fully-imaged level representations are corrected via sphering transformation (Fig. 5b). Finally, the fully-imaged level representations are aggregated to treatment level representations by computing their mean (Fig. 5c).

**Fig. 5.**
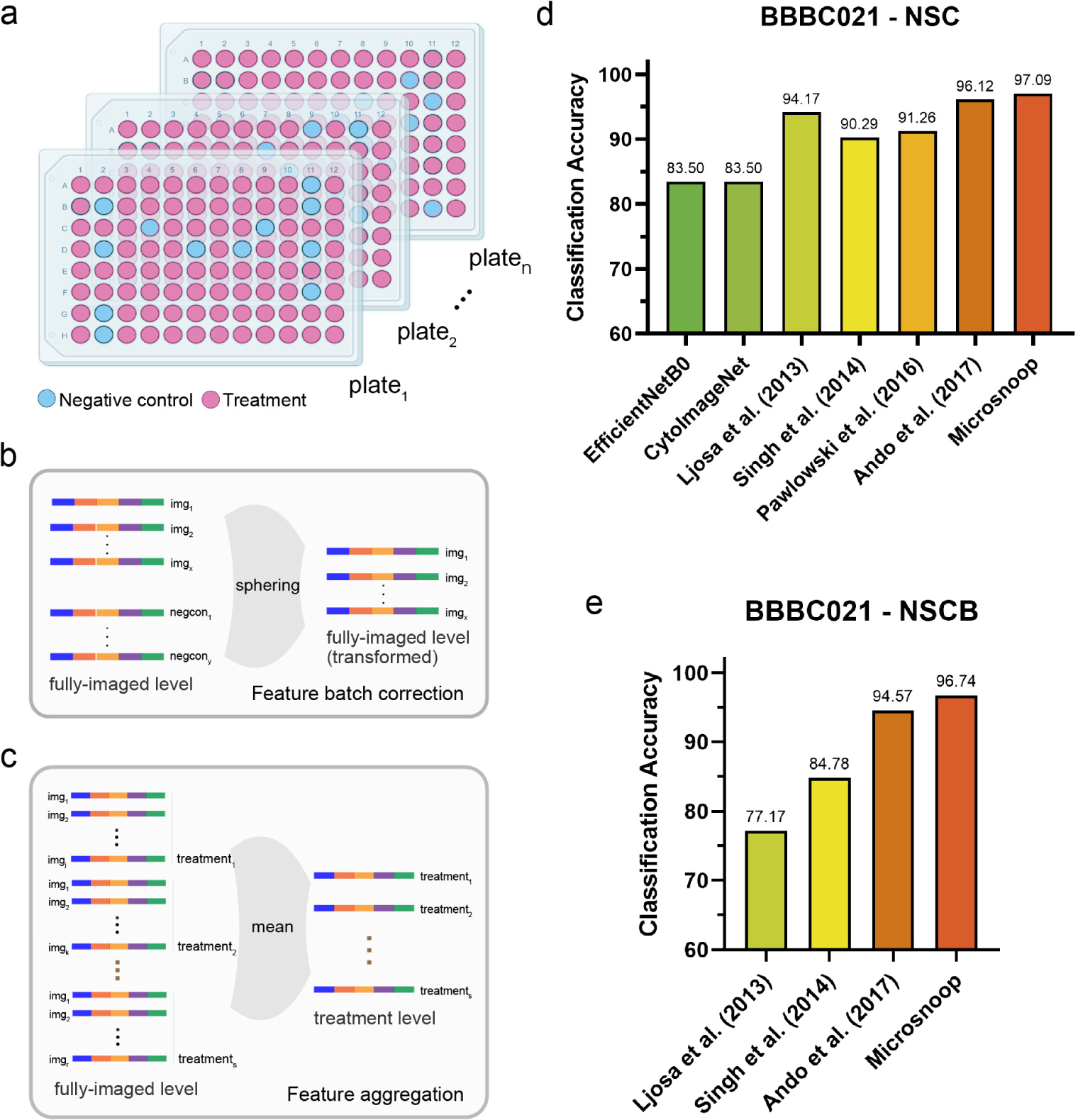
Profiling with Microsnoop on batch-experiment images. **a,** Schematic of multi-well plates in a drug screening experiment containing negative control wells and different treatment wells set in each plate. **b,** Batch correction on fully-imaged level representations. **c,** Feature aggregation on fully-imaged level embeddings to obtain treatment level image representations. **d,e,** Benchmark on BBBC021, with different evaluation metrics.

We evaluated the representation ability of Microsnoop on the classic BBBC021 dataset, while including previously reported results of generalist and custom methods in the comparisons. We assessed the performance with the Not-Same-Compound (NSC) and Not-Same-Compound-or-Batch (NSCB) KNN classification accuracy. Microsnoop still achieved state-of-the-art performance without using any data from the dataset, even if compared with the methods exclusively studied on it (Fig. 5d-e).

### Two other fully-imaged image profile modes and the robustness of cell region cropping mode

In addition to the cell region cropping mode, we provided two alternative modes for processing fully-imaged datasets: rescaling and tile mode. In the rescaling mode, the shape of the fully-imaged images is directly rescaled to the input size (224*224) as inputs (Extended Data Fig. 6a-b). In the tile mode, the original image is cropped into multiple 224×224 tiles, and the fully-imaged level representations are aggregated by computing the mean over all tiles (Extended Data Fig. 6c). We evaluated the performance of these three processing modes, including different rescale constants for the cell region cropping mode, on both the fully-imaged and batch-experiment datasets (Extended Data Fig. 6d-g and Methods). The rescaling and tile modes outperformed the single-cell mode on LIVECell and TissueNet tests; however, both modes displayed a significant performance decline on the BBBC021 dataset. The reason for the underperformance of the rescaling mode could be attributed to the fact that it discards high-resolution phenotypic information during the rescaling process. On the other hand, the decline in performance observed with the tile mode may be due to the fact that it averages out important subtle phenotype variations present in certain regions of fully-imaged images. In contrast, the cell region cropping mode displayed robust performance across a range of parameters on all three datasets. Although the single-cell mode is more robust and efficient, it requires more time and memory compared to the other two modes. (Extended Data Fig. 6h-i).

### Microsnoop improves the performance of the multi-modal structured embedding algorithm

A recent study of the multi-modal structured embedding algorithm (MUSE^17^) has shown impressive results for the integrative spatial analysis of image and transcriptional data. The authors conducted a simulation experiment to assess the performance of MUSE when transcriptional data quality is degraded. Here, we focused on the impact of image feature quality, and the results of our simulation experiment showed that with the quality improvement of image representations, the performance of MUSE can also be significantly improved (Extended Data Fig. 7). Next, we tested Microsnoop on the real-world dataset seqFISH+^8^ in comparison with the representation method used in the original paper. After acquiring the image representations, we use principal component analysis (PCA) performing feature dimensionality reduction to match the latent space dimensions of MUSE (Fig. 6a). We employed the silhouette coefficient^48^ to evaluate the feature quality. Microsnoop demonstrated better image representation quality and greater improvement in the performance of MUSE (Fig. 6b).

**Fig. 6.**
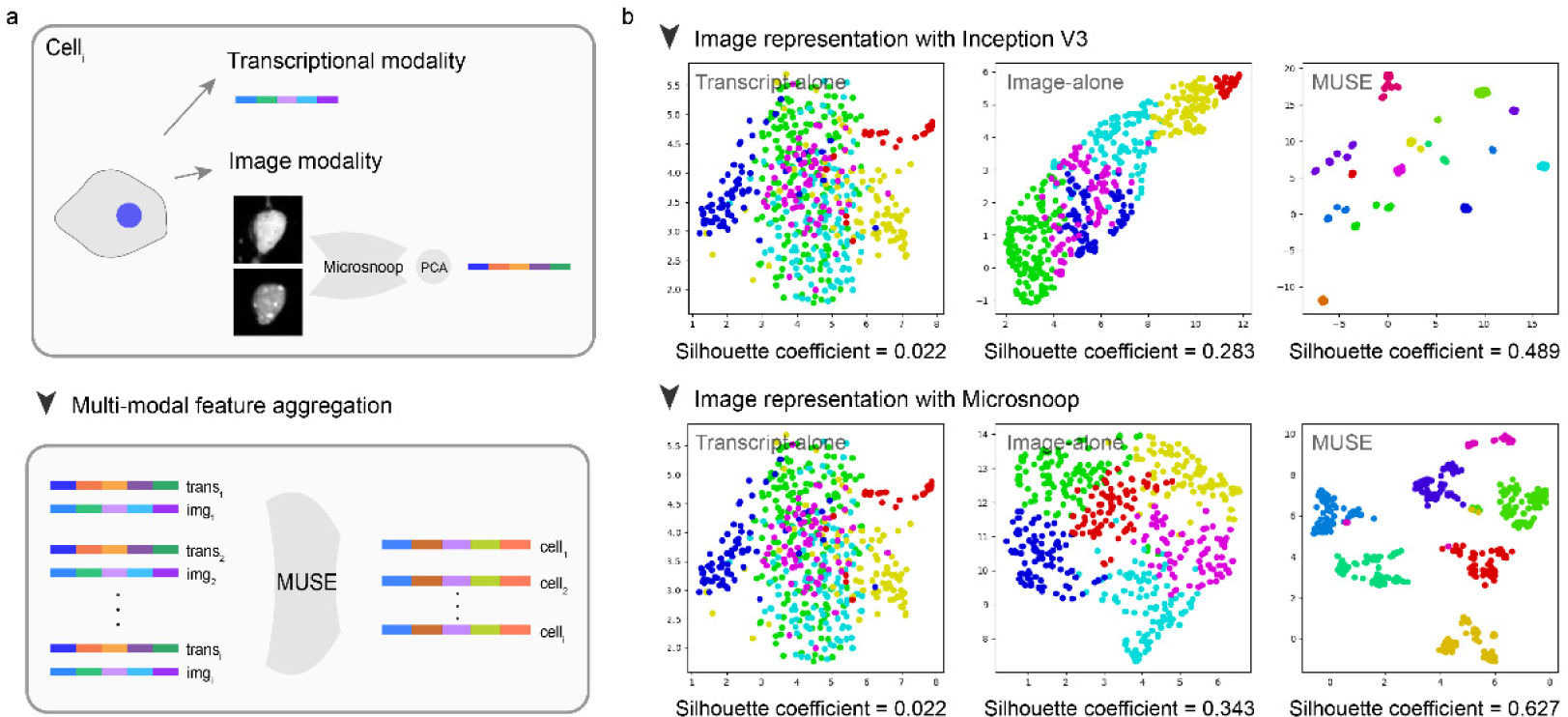
Joint use of Microsnoop and MUSE. **a,** Pipeline. Image modality data is first processed by Microsnoop, then PCA is performed on the output representations to reduce feature dimensionality. Finally, two modality representations are mixed by MUSE. **b,** UMAP visualization of different modality latent spaces on seqFISH+, using two different image representation methods. Silhouette score was used to quantify the separateness of clusters.

## Discussion

Advances in imaging technology, such as phase-contrast microscopy, imaging flow cytometry, automated high-throughput microscopy and microscopy combined with spatial omics techniques have created a massive demand to solve the complex challenge of microscopy image representation. In this study, we present Microsnoop, an innovative deep learning tool that effectively addresses this challenge. The accurate analysis of heterogeneous microscopy images, as a critical aspect of both fundamental and applied biological research, is highly valued by the microscopy image analysis community^49–50^. Our proposed solution offers promising advancements to this field. Microsnoop was trained on large-scale high-quality data using a masked self-supervised pretext task, allowing it to learn valuable and unbiased features for image representation. The one-channel feature concatenation strategy, efficient task distribution module, and rational classification mode of profiling needs make our tool flexible to meet various user needs. In addition, Microsnoop is capable of processing complex fully-imaged images through cell region cropping and mitigating batch effects in batch-experiment images through sphering transformation. For fully-imaged images, our results show that the single-cell analysis mode is more robust compared to other modes, reinstating the importance of considering cellular heterogeneity in biological research. Our benchmark results demonstrate robust and state-of-the-art performance on all evaluated datasets, eliminating the need to use of any evaluation data for fine-tuning. Furthermore, the enhanced representation of unimodal image data leads to significant improvements in the performance of multi-modal algorithms.

In our methodology experiments, we found that a mask ratio of 25% is optimal for microscopy images, which is significantly lower than the 75% that has been found optimal for natural images^21^. The difference is primarily due to the smaller size and erratic content of instances in microscopy images, which may result in lost information if too much reference information is masked. Compared with the CytoImageNet^18^ study that utilized a supervised classification task as the pretext task, our masked self-supervised learning approach only requires raw images without any manual annotation and yields unbiased and more capable representations. Recently, a similar self-supervised representation learning study has also been reported as useful in learning the representations of protein subcellular location images through a pretext task that requires the network to directly reconstruct original images and images corresponding to similar proteins having similar representations^19^. In contrast, the uniqueness of our method is that ours do not require domain-specific knowledge and is developed for generalist image representation. Our benchmark study has shown that a single network is capable of handling heterogeneous microscopy images, which is in line with the conclusion reached in the sister domain of cell segmentation^25^. Furthermore, our pretext task was trained on the same network structure as Cellpose. This is reminiscent of the recent success of large pre-trained language models in the field of natural language processing^51–53^. With continued advancements in the understanding of computer vision and the further development of models for microscopy image representation and other image processing tasks, such as cell segmentation, it may be possible to merge these models into a single, unified model in the future.

While Microsnoop is a powerful tool, there are several areas for improvement. For example, further evaluation is needed to determine the efficacy of our approach of one-channel feature concatenation and feature aggregation in 3D and time-series imaging datasets in comparison to training a network to directly extract spatial or temporal information. To enhance the capabilities of Microsnoop, future work could include incorporating additional self-supervised pretext tasks for multi-task learning, optimizing the quality of the training dataset and refining the single-cell level feature aggregation methods. Moreover, the current training images are still limited in size compared to natural images, and a larger training data volume combined with the Transformer architecture can be studied to improve the performance. Last but not least, deploying our model on mobile devices to aid rapid detection could be a valuable application scenario^54^.

Overall, we have developed an impressive, generalist tool for microscopy image representation. We anticipate its positive impact on the microscopy image analysis community, facilitating new phenotype discovery, data sharing, and the establishment of large image databases, among other benefits. Furthermore, we envision that Microsnoop can be effectively utilized in multi-modal studies such as combining molecular and image representation for MoA prediction^55–56^ or exploring the relationship between gene expression, image representation for drug discovery^57^ and much broader applications^58–59^.

## Methods

### Training set

The training set consisted of four diverse image types from seven published datasets: Cellpose, LIVECell, TissueNet, and Histo, which includes MoNuSeg, MoNuSAC, and NuCLS. Firstly, all channels of the images were separated. For Cellpose and TissueNet, only the cell body channel was utilized, while the original RGB images of Histo were transformed into grayscale. The original training-validation dataset split was maintained for Cellpose, LIVECell, and TissueNet, while the images from the three Histo subsets were mixed and 20% were randomly reserved for validation purposes. Finally, the training set was organized into a one-channel image data pool. A comprehensive summary of the training set can be found in Supplementary Table 1.

### Model architecture

The network architecture was based on a refined version of the classic U-Net^34^, as utilized in Cellpose. The standard convolutional blocks were replaced with residual blocks and style embeddings were incorporated into the concatenation stages. The downsampling scale was set as 32, 64, 128 and 256, and the upsampling scale was mirror symmetry. Both the input and output tensors were of shape batch_size*1*224*224 (in Pytorch tensor format, where batch_size is described below).

### Masked self-supervised learning

In the masked self-supervised learning approach, the network is tasked with reconstructing the original image from partial masked images. Our implementation involved dividing the target image (after normalization and augmentation) into 16*16 non-overlapping patches. Subsequently, a portion of these patches were randomly replaced with black patches of size 16*16, where every pixel was zero. Different from the original MAE built on a Transformer architecture, the transformed patches were restored to the image format to accommodate the input format of the CNN architecture.

### Model training

The self-supervision loss was set as the mean square error loss (MSE), which calculates the difference in both the masked and unmasked areas. The network was optimized by AdamW optimizer from the torch.optim Python package. In our implementation, we adopted a different definition of an epoch, in which one epoch corresponds to a complete iteration through all the sampled data, rather than through all the training data, as is commonly defined. During each epoch, we randomly sampled 12000 images from the four different types of training data in turn. The batch size was set as 16. The initial learning rate was set as 0.001, and we used a learning rate (LR) warmup trick: at the first 40 epochs, the LR was computed as:

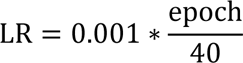

after 40 epochs, the LR was computed as:

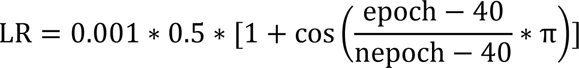

where nepoch represents the epoch size of the training process, here it was set as 1000.

### One-channel feature concatenation strategy for multi-channel image representation

In our implementation of Microsnoop for feature extraction, we assumed that the input data comprised multi-channel images with the same number of channels, represented as (c, h, w), where c denotes the number of channels, and h and w denote the height and width, respectively. In the event that images had different h and w, we padded them with zeros to obtain a consistent shape. The task distribution module is then used to read the images into CPU memory, where they are transformed into an array with shape (n, c, h, w), where n denotes the number of images read. This array is then reshaped into (n*c, 1, h, w), with each image assigned a unique index represented as a shape (n*c,) vector. For each batch of size b, the task distribution module transfers b images into the GPU memory, resulting in a tensor of shape (b, 1, h, w). After Microsnoop processes all n*c images, the CPU cache is cleared using the collect function from the gc Python package, and the next n images are read. The resulting embedding array had the shape of (N*c, 256), where N denotes the total number of processed images, and 256 is the pre-set dimensionality of the feature vector for a one-channel image in Microsnoop. These embeddings are then concatenated in channel to obtain a final feature embedding array of shape (N, 256*c).

### Evaluation datasets

We curated seven evaluation datasets, four of which were directly available from public sources and three (CoNSeP, LIVECell Test and TissueNet Test) were processed by us based on publicly acquired images. The summary of these datasets can be seen in Supplementary Table 2.

#### COOS7

This dataset contains 132,209 single-cell fluorescence images, including a training set and four test sets that vary in different factors. The training set consists of images from 4 independent plates, while Test 1 includes randomly held-out images from the same plates as the training set, Test 2 includes images from the same plates but different wells, Test3 comprises images produced months later, and Test 4 has images produced by other instruments. The images were downloaded through the link provided by Stanley Bryan Z. Hua^18^. Each image takes the shape of 2*64*64 and is a pixel crop centered around a unique mouse cell. One channel marks the protein targeting a specific component of the cell and the other marks the nucleus. There are 7 protein location classes in each set: Endoplasmic Reticulum, Inner Mitochondrial Membrane, Golgi, Peroxisomes, Early Endosome, Cytosol and Nuclear Envelope, and the evaluation task requires the model to accurately predict the protein location.

#### CYCLoPs

This dataset consists of 28,166 single-cell fluorescence images from the CYCLoPs database, and we downloaded the data through the link provided by Stanley Bryan Z. Hua^18^. Each image has a shape of 2*64*64 and is a pixel crop centered around a unique yeast cell. One channel marks the protein location and the other marks the cytosol. There are 17 protein location classes: ACTIN, BUDNECK, BUDTIP, CELLPERIPHERY, CYTOPLASM, ENDOSOME, ER, GOLGI, MITOCHONDRIA, NUCLEARPERIPHERY, NUCLEI, NUCLEOLUS, PEROXISOME, SPINDLE, SPINDLEPOLE, VACUOLARMEMBRANE and VACUOLE. The aim of the evaluation is to accurately predict the protein localization.

#### CoNSeP

This dataset has 41 H&E stained fully-imaged images with a shape of 3*1000*1000 pixels. 14 of these are test images and 27 are training images. The raw data were obtained from https://warwick.ac.uk/fac/sci/dcs/research/tia/data and then transformed into grayscale format. Each cell was cropped based on the provided segmentation mask, resulting in 8777 single-cell test images and 15554 single-cell training images with a shape of 1*112*112 pixels. In cases where the cells were smaller, padding was applied to obtain the desired size. The class information was extracted from the classification mask, with 4 classes: Other, Inflammatory, Epithelial, Spindle-shaped. The evaluation task requires the model to accurately predict the cell types.

#### BBBC048

This dataset contains 32,266 single-cell images from the Broad Bioimage Benchmark Collection^60^. These single-cell images of Jurkat cells were directly captured with the ImageStream imaging flow cytometer. Each image has a shape of 3*66*66 pixels, with a brightfield channel and two fluorescence channels. There are 7 cell phases: G1, S, G2, Prophase, Metaphase, Anaphase and Telophase. Another 5-phase case considers G1, S and G2 phase as a single class. The evaluation task requires the model to accurately predict the cell cycle stages.

#### LIVECell Test

This dataset comprises 1512 fully-imaged phase-contrast images provided by Christoffer Edlund^26^, where each image has a shape of 1*520*704 pixels. There are 8 cell types: A172, BT474, BV2, Huh7, MCF7, SHSY5Y, SkBr3 and SKOV3. The evaluation task requires the model to accurately predict the cell types of full-imaged images.

#### TissueNet Test

This dataset comprises 1249 fully-imaged tissue images provided by Noah F. Greenwald. Each image has a shape of 2*256*256 pixels, one channel marks the membrane or cytoplasm and the other marks the nucleus. We extracted the tissue type information from the metadata provided. There are 6 tissue types: Breast, Gi, Immune, Lung, Pancreas and Skin. The evaluation task requires the model to accurately predict the tissue types of full-imaged images.

#### BBBC021

This dataset includes 3848 fully-imaged fluorescence images, a subset from the Broad Bioimage Benchmark Collection^60^. The images are of MCF-7 breast cancer cells with a collection of 113 small molecules at different concentrations and a DMSO negative control. Each image has a shape of 3*1024*1280 pixels, and different channels respectively mark the DNA, F-actin and B-tubulin. There are 12 mechanisms: Actin disruptors, Aurora kinase inhibitors, Cholesterol-lowering, DNA damage, DNA replication, Eg5 inhibitors, Epithelial, Kinase inhibitors, Microtubule destabilizers, Microtubule stabilizers, Protein degradation and Protein synthesis. The evaluation task requires the model to accurately predict the MoA of different treatments.

### Three modes for the profile of fully-imaged images

#### Cell region cropping mode

We utilized the generalist tool Cellpose on the easiest channel (such as the nucleus channel) to perform cell segmentation. For each image, following the acquisition of the segmentation mask, we extract all the (x, y) pixel coordinates of each cell, and compute the region of each cell as follows:

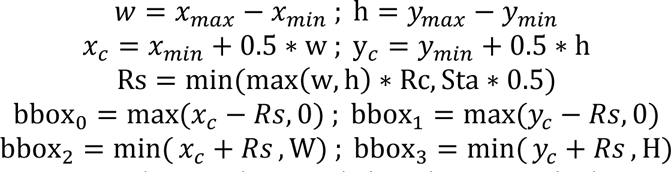

where *x_max_*, *x_min_*, *y_max_*, *y_min_* denote the max/min x/y, respectively, among all the pixels coordinates; *x*_c_, *y*_c_ denote the coordinates of centroid; Rc denotes the rescale constant (it is set by user according to the average size of cell bodies); Sta denotes the side length of cropped image (here we set it as 224, the input size of Microsnoop); Rs denotes the crop size (it cannot be more than half of Sta); W, H denote the width and height of the fully-imaged image, respectively. bbox_o_, bbox_1_, bbox_2_, bbox_3_ denote the left, up, right, down of the cropped region in the original image, respectively, and they cannot go beyond the boundaries of the image. Finally, single-cell images are cropped on all channels and padded to (c, Sta, Sta) with zero pixels if smaller, where c denotes the number of channels. The fully-imaged level embedding of the image is obtained by computing the mean of all single-cell image embeddings.

#### Rescaling mode

In the case that the height of the image is not equal to its width, the initial step is to pad the image with zeros to create a square shape. The fully-imaged images are then rescaled to input size using the resize function from the cv2 Python package. The fully-imaged level embedding of the image is directly obtained through this process.

#### Tile mode

The fully-imaged images are cropped into tiles using the make_tiles function from the cellpose.transforms Python package. The parameter bsize was set as the input size, and the parameter tile_overlap was set as 0.1. The fully-imaged level embedding of the image is obtained by computing the mean of all tile embeddings.

### Sphering transformation for the profile of batch-experiment images

The detailed description can be found in ref. ^47^. Here, we fitted the ZCA_corr transformer from https://github.com/jump-cellpainting/2021_Chandrasekaran_submitted/blob/main/benchmark/old_notebooks/utils.py on the embeddings of negative control, and then used the fitted transformer to correct the embeddings of each batch.

### Benchmarking

For BBBC021, we directly adopted the previously published state-of-the-art (SOTA) results from the curated resource at https://bbbc.broadinstitute.org/BBBC021. We also included the results of recently reported generalist methods. All results were formatted to two decimal places.

For other datasets, we compared with three generalist deep-learning methods: EfficiententNetB0, Inception V3 and CytoImageNet. EfficiententNetB0 was pretrained on the ImageNet and was included in the comparison in CytoImageNet. The famous project DeepProfiler^47^ also used this network for the profiling of microscopy imaging data. Inception V3, which was also pre-trained on ImageNet, had been utilized in the MUSE project, a study of advanced multimodal algorithms. CytoImageNet, a recently published generalist microscopy image representation learning algorithm, was pre-trained using a self-constructed microscopy image classification dataset.

The results of EfficiententNetB0 and CytoImageNet on COOS7 and CYCLoPs have been previously reported^18^ and were directly adopted from the relevant publication. For BBBC048, we also included the custom algorithm results reported in the original paper. The remaining results presented in this paper were generated by the authors.

EfficiententNetB0 and CytoImageNet were established using the EfficientNetB0 class from the tenforflow.keras.applications Python package, with different weights loaded (EfficiententNetB0 used the ImageNet weights and CytoImageNet used the weights published by Stanley Bryan Z. Hua). Inception V3 was established using inception_v3 class from the torchvision.models Python package. We dropped the last classification layer and used the remaining network for feature extraction. Because these network architectures are presented in natural RGB image study, at test time, each one-channel image is copied three times to mimic RGB images (also used in ref. ^18, 37^). The other steps, such as data preprocessing and feature aggregation, are identical to those used in the Microsnoop protocol.

For LIVECell and TissueNet Test, we directly used the provided segmentation masks (nucleus channel for the TissueNet) without applying the cell segmentation algorithm in the cell region cropping mode. For the COOS7, CYCLoPs and BBBC021 datasets, the number of nearest neighbors (k) in the KNN classifier was set to 11, 11, and 1, respectively, in accordance with the ref. ^18^. For BBBC048, the MLP was conducted using the MLPClassifier class from the sklearn.neural_network Python package, and the parameter max_iter was set as 1000.

### Joint use of Microsnoop and MUSE

In the simulation experiment, we utilized the simulation_tool.multi_modal_simulator function from the MUSE project to generate the transcriptional and image representations along with the corresponding ground truth. We used the adjusted Rand index (ARI)^61^ to assess the ability of discovering true subpopulations. For the analysis of seqFISH+ data, the microscopy images were provided by the authors of the seqFISH+ paper. Each cell region of the images was determined by the coordinates of the cell centroid provided. We used Microsnoop and Inception V3 to conduct feature extraction on the Nissl and DAPI stained images separately. The shape of each single-cell embedding output was 512 (256*2), then we used PCA to reduce the feature dimensionality to 500. The process of the transcript data was the same as MUSE. We used the silhouette coefficient to assess feature quality by the compactness of the clusters, which was conducted using the silhouette_score function from the sklearn.metrics Python package.

## Graph plotting

All bar graphs were plotted using GraphPad PRISM 8.0 software (GraphPad Software, Inc., CA, USA). Fig. 1b(i) and Fig. 5a were created using resources from BioRender.com. The sources of images in Fig. 1 also included https://www.rxrx.ai/rxrx2, in addition to those listed in the supplementary Table 1 & 2. Some microscopy images in the figures have been processed using “Enhance Contrast…” from ImageJ^62^ for better presentation.

## Software and hardware

The programming was conducted using Python v.3.7. Training and all evaluations were performed on NVIDIA GeForce RTX 3090 GPUs. The deep learning framework of Microsnoop used PyTorch^63^ v.1.10.

## Data availability

The links to download the raw data of training set and evaluation datasets are provided in Supplementary Table 1-2. The new evaluation datasets generated by this study are available on figshare: https://figshare.com/articles/dataset/Microsnoop_a_generalist_tool_for_the_unbiased_representation_of_heterogeneous_microscopy_images/22197607.

SeqFISH+ mouse cortex dataset: Transcript data were downloaded from https://github.com/CaiGroup/seqFISH-PLUS. Image data were provided by the authors of the seqFISH+ paper.

All data in this study are available from the corresponding author upon reasonable request.

## Code availability

Source code for Microsnoop, including detailed tutorial, is available on GitHub (https://github.com/cellimnet/microsnoop-publish). A configured Amazon Machine Image (AMI) will be made available upon publication for quickly and conveniently deploying Microsnoop for microscopy image analysis.

## Acknowledgments

We thank L. Cai at Caltech for providing seqFISH+ image data. We thank W.K. Wang and L. Sun at Amazon Web Services China for their indispensable support in terms of computing resources and technology. We are grateful for the support from ZJU PII-Molecular Devices Joint Laboratory and support from “Medicine + X” interdisciplinary Center of Zhejiang University. This study was supported by National Key R&D Program of China (2021YFC1712905), National Natural Science Foundation of China (No. 82173941, No. 61872319), Key R&D Program of Zhejiang Province (No. 2023C01039). Y.W. was supported by the Innovation Team and Talents Cultivation Program of National Administration of Traditional Chinese Medicine (No. ZYYCXTD-D-202002).

## Author contributions

Y.W., X.C.Z. and R.W. supervised the study, D.J.X. acquired data, established pipeline, conducted experiments and performed data analysis. D.J.X., Y.W., X.C.Z. and R.W. wrote the manuscript.

## Extended Data

**Extended Data Fig. 1.**
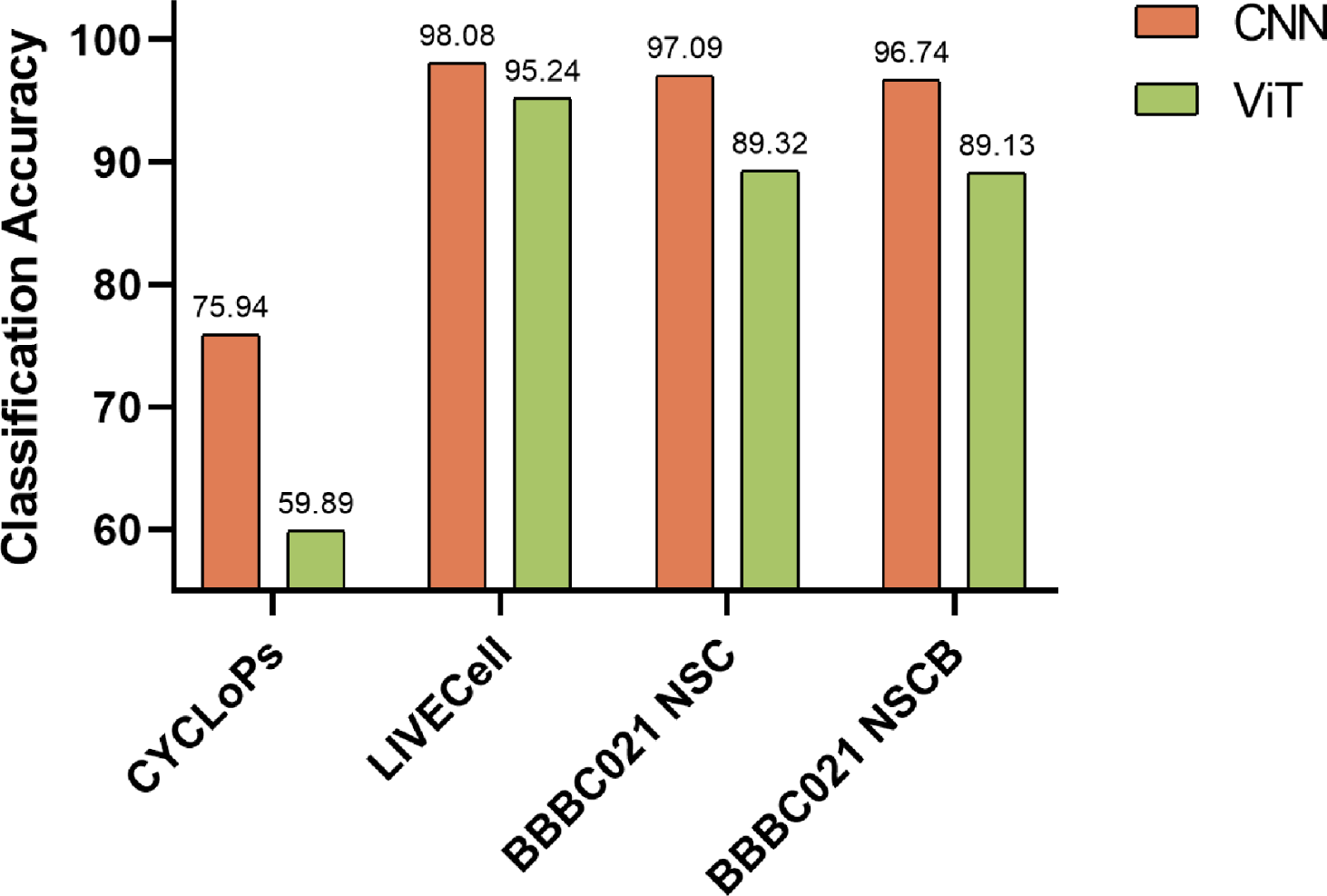
Performance evaluation of Microsnoop trained with different network architectures. Three representative datasets from seven evaluation datasets were selected for the early trials: single-cell image task (CYCLoPs), fully-imaged image task (LIVECell), and batch-experiment image task (BBBC021). The ViT architecture referred to the MAE, and the classification accuracy for the corresponding dataset was reported.

**Extended Data Fig. 2.**
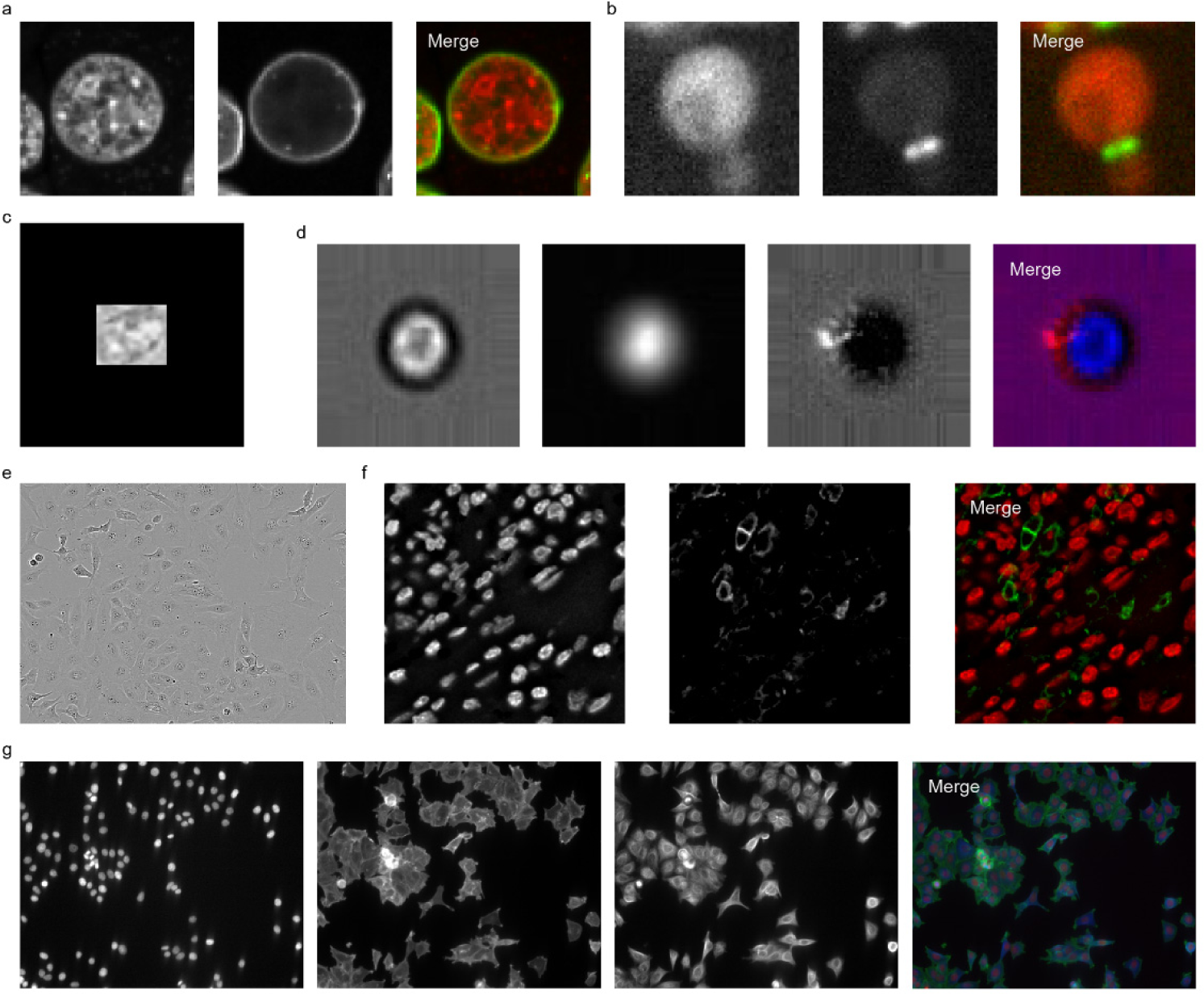
Example images of evaluation datasets. Each channel of the example image was presented for each dataset: **a,** COOS7 **b,** CYCLoPs **c,** CoNSeP **d,** BBBC048 **e,** LIVECell **f,** TissueNet **g,** BBBC021.

**Extended Data Fig. 3.**
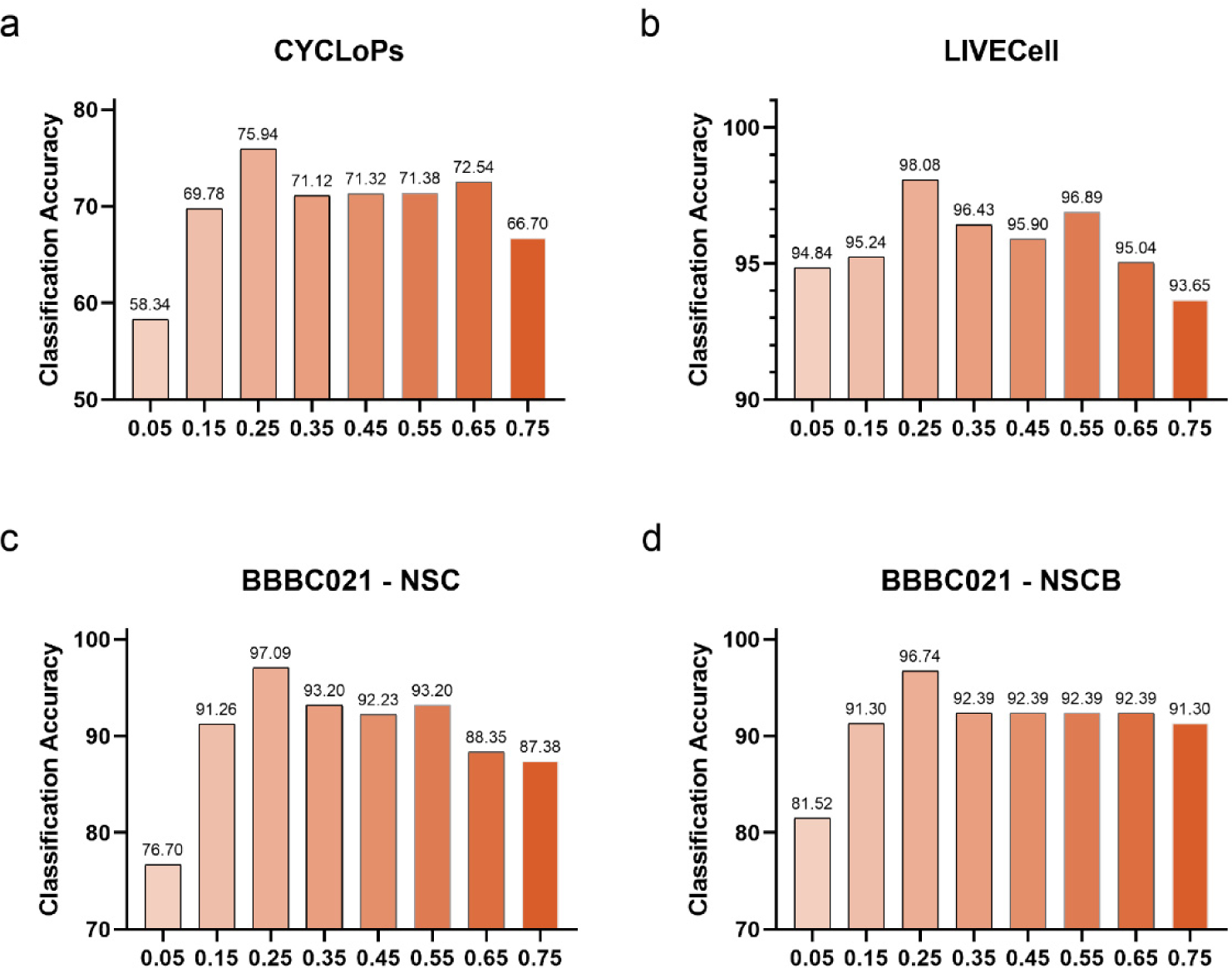
Performance evaluation of Microsnoop trained with different mask ratios. Three representative datasets from seven evaluation datasets were selected for the early trials: **a,** Single-cell image task **b,** Fully-imaged image task **c,d,** Batch-experiment image task. The mask ratio was set ranging from 0.05 to 0.75, and the classification accuracy for the corresponding dataset was reported.

**Extended Data Fig. 4.**
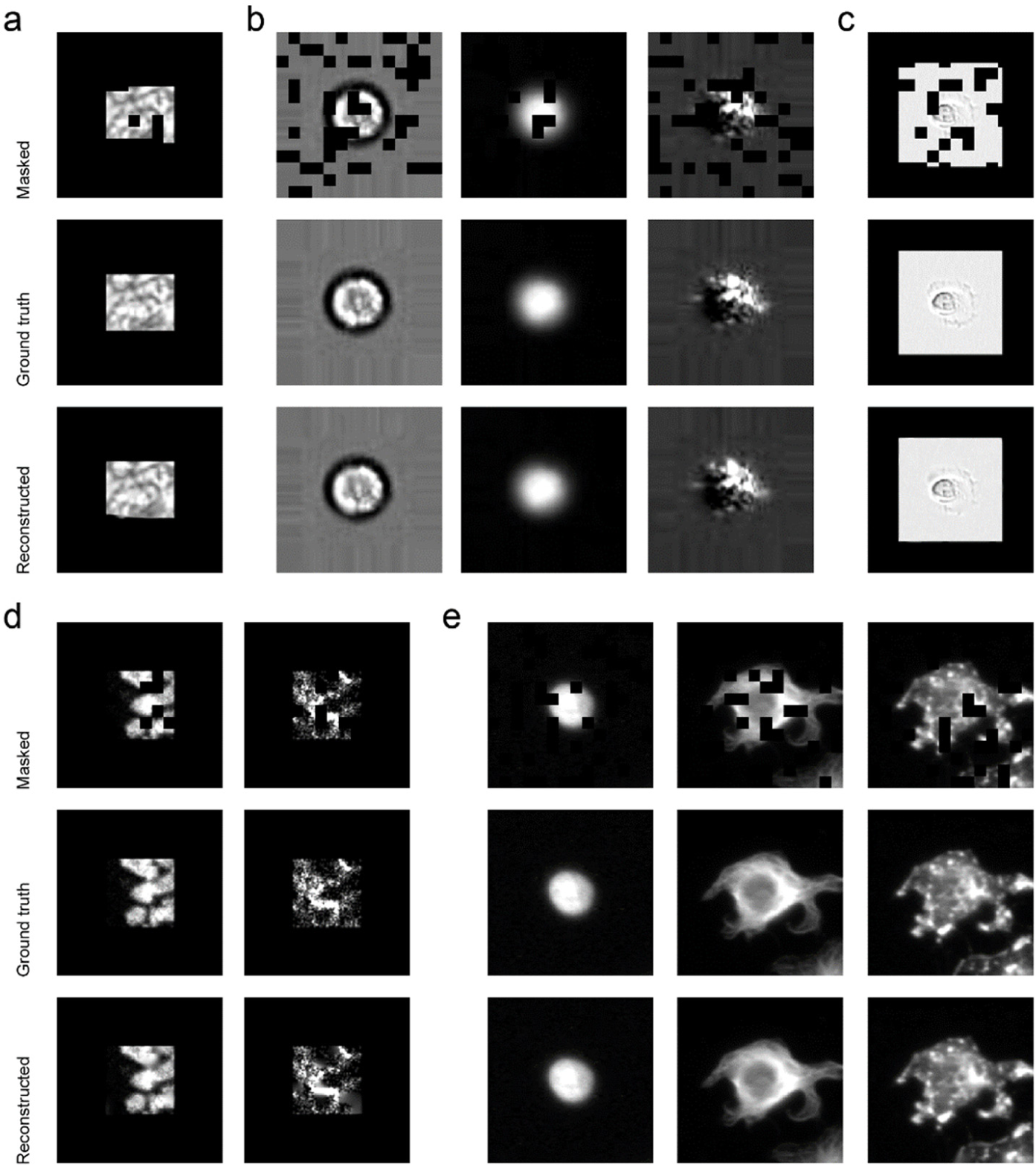
Reconstruction results with Microsnoop on the remaining evaluation datasets. Each channel of the example images from each dataset were performed: **a,** CoNSeP **b,** BBBC048 **c,** LIVECell **d,** TissueNet **e,** BBBC021. For fully-imaged image datasets (**c-e**), the processed single-cell images after cell region cropping were used.

**Extended Data Fig. 5.**
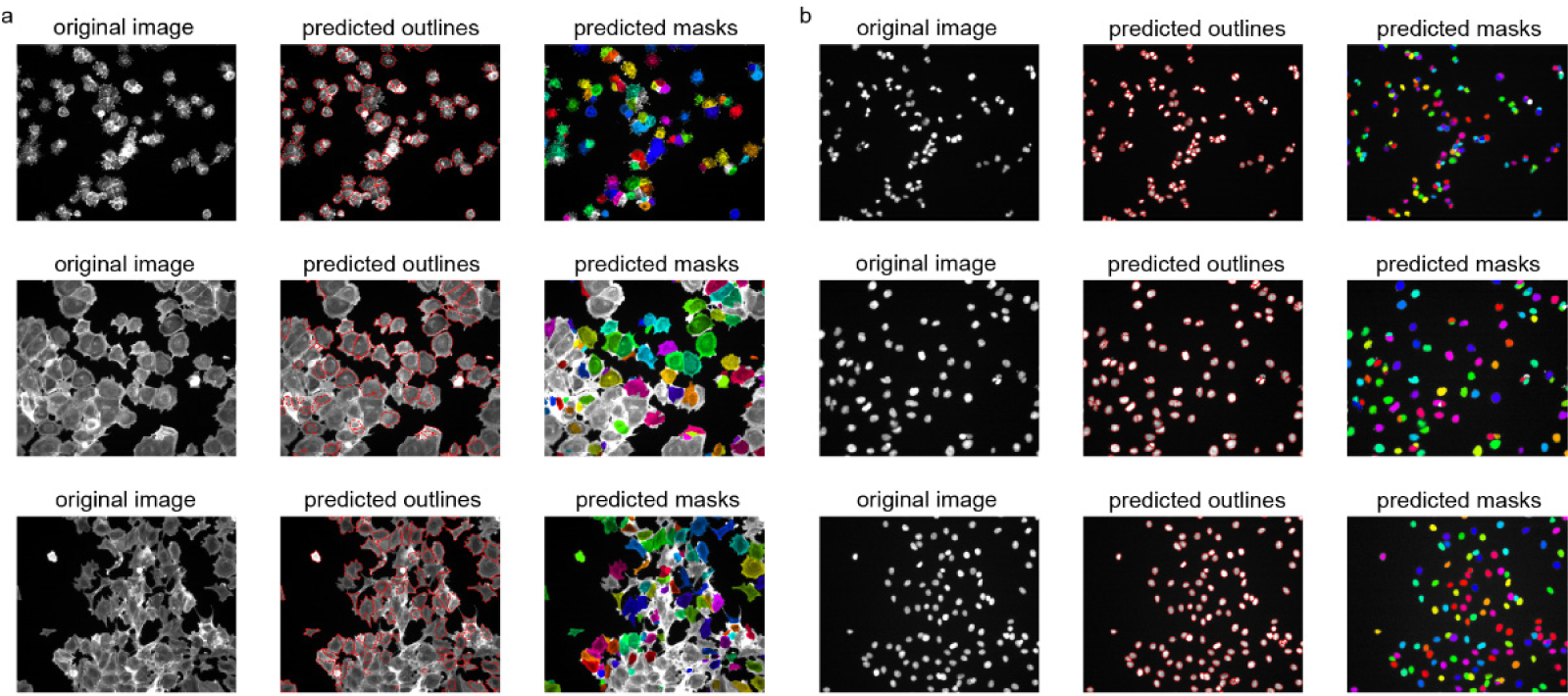
Example segmentation results of the generalist model for high-content screening images. Images were shown in pairs, with the original image on the left and the segmentation results on the right using two visualization methods; the predicted outlines show the boundary of each cell and the predicted masks mark the segmented cells with different colors. Three images were selected from the BBBC021 dataset, in which cells were treated with different compounds and presented complex phenotypes. Cell segmentation was conducted with Cellpose. **a,** Segmentation on F-actin channel images. **b,** Segmentation on corresponding nucleus channel images.

**Extended Data Fig. 6.**
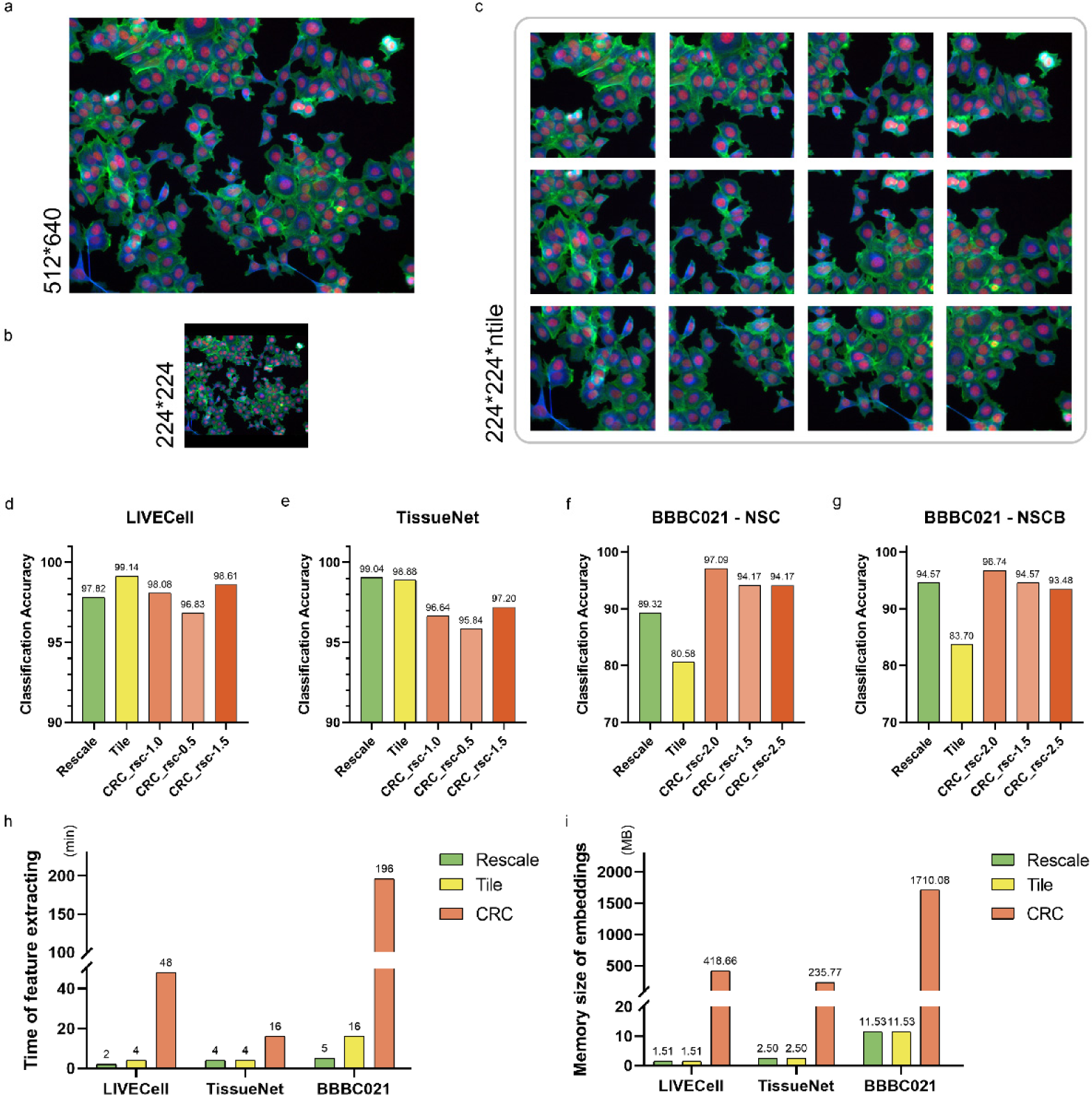
Different profile modes of fully-imaged images. **a,** An example image. **b,** Example of the rescaling mode, where the original image was patched and rescaled to the input size (224*224). **c,** Example of the tile mode, where the original image is cropped to many 224*224 tiles (ntile) using the make_tiles function from the cellpose.transforms Python package, and the tile_overlap parameter was set as 0.1. **d-g,** Performance comparison of different modes on three evaluation datasets: **d,** LIVECell **e,** TissueNet **f,g,** BBBC021. The cell region cropping mode (CRC) was tested with different rescale constant to study the robustness. **h,i,** Time (**h**) and memory (**i**) cost of different modes. In the case of CRC mode, the memory cost computes the representations of all single-cell images, rather than the final fully-imaged level image representation.

**Extended Data Fig. 7.**
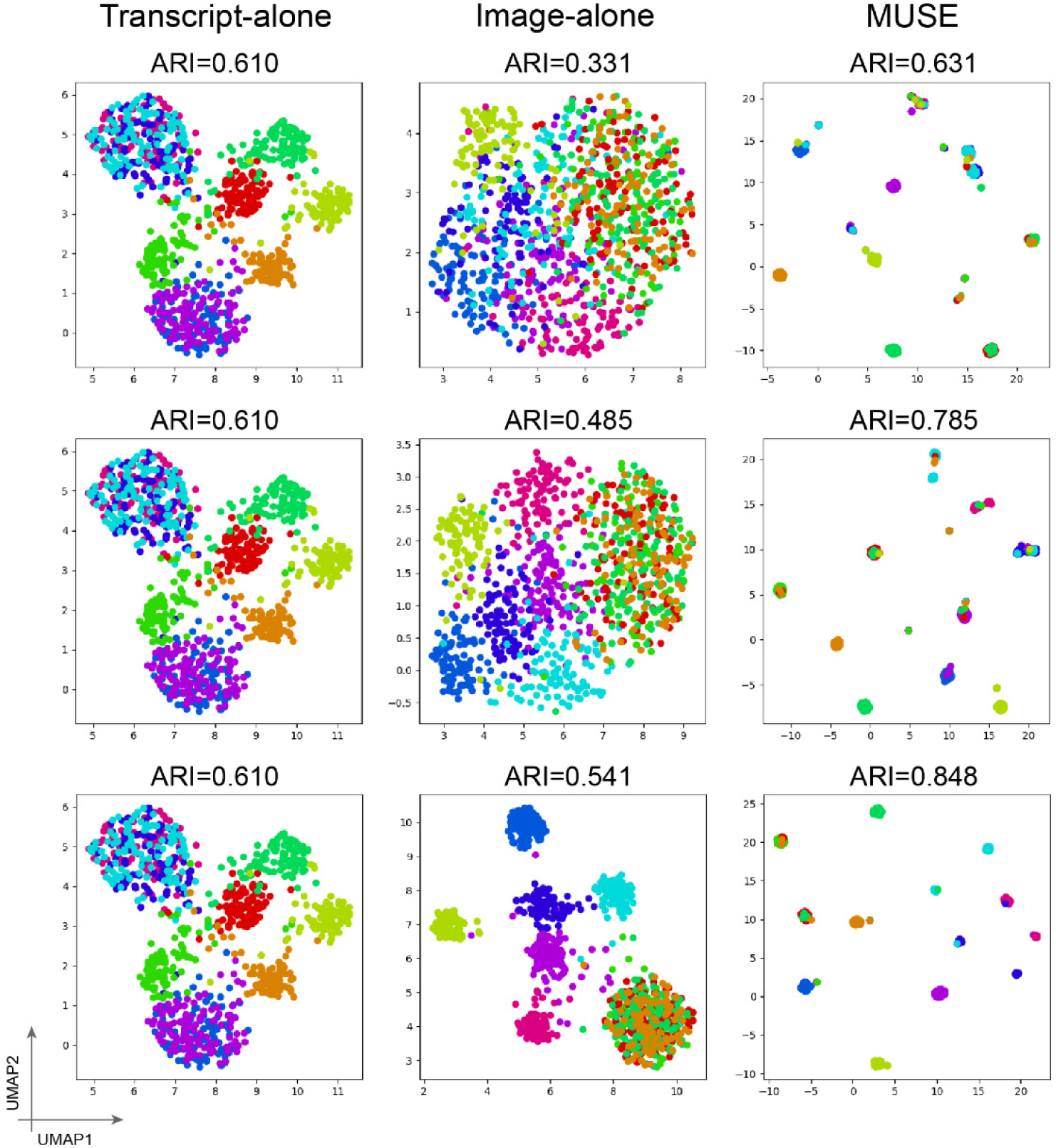
UMAP visualizations of latent embeddings from single- and combined-modality methods. Colors: ground truth subpopulation labels in simulation. Cluster accuracy is quantified using the adjusted Rand index (ARI).

